# HIV-1 envelope glycoproteins proteolytic cleavage protects infected cells from ADCC mediated by plasma from infected individuals

**DOI:** 10.1101/2021.10.26.465908

**Authors:** Jérémie Prévost, Halima Medjahed, Dani Vézina, Hung-Ching Chen, Beatrice H Hahn, Amos B. Smith, Andrés Finzi

## Abstract

The HIV-1 envelope glycoprotein (Env) is synthesized in the endoplasmic reticulum as a trimeric gp160 precursor, which requires proteolytic cleavage by a cellular furin protease to mediate virus-cell fusion. Env is conformationally flexible, but controls its transition from the unbound “closed” conformation (State 1) to downstream CD4-bound conformations (States 2/3), which are required for fusion. In particular, HIV-1 has evolved several mechanisms that reduce the premature “opening” of Env which exposes highly conserved epitopes recognized by non-neutralizing antibodies (nnAbs) capable of mediating antibody-dependent cellular cytotoxicity (ADCC). Env cleavage decreases its conformational transitions favoring the adoption of the “closed” conformation. Here we altered the gp160 furin cleavage site to impair Env cleavage and to examine its impact on ADCC responses mediated by plasma from HIV-1-infected individuals. We found that infected primary CD4+ T cells expressing uncleaved, but not wildtype, Env are efficiently recognized by nnAbs and become highly susceptible to ADCC responses mediated by plasma from HIV-1-infected individuals. Thus, HIV-1 limits the exposure of uncleaved Env at the surface of HIV-1-infected cells at least in part to escape ADCC responses.

## 1. Introduction

The human immunodeficiency virus type 1 (HIV-1) envelope glycoprotein (Env) is a class I viral membrane fusion protein which mediates viral entry using the CD4 cellular receptor. The envelope gp160 precursor is synthesized in the endoplasmic reticulum (ER) and oligomerizes as a trimer [1, 2]. Subsequently, the trimeric Env traffics through the trans-Golgi network (TGN) to reach the plasma membrane and to be incorporated into nascent HIV-1 virions [3–5]. During its transit through the secretory pathway, Env undergoes important post-translational modifications, including N-linked and O-linked glycosylation as well as proteolytic cleavage [6–10]. The addition of high-mannose oligosaccharides takes place in the ER and these glycans are further processed to acquire complex modifications in the TGN [11]. Concomitantly, proprotein convertases present in the TGN, including furin and furin-like proteases, catalyze the cleavage of the immature gp160 polyprotein [12–15] into two functional non-covalently linked subunits: the exterior gp120 subunit, which is responsible for viral attachment and the transmembrane gp41 subunit, which mediates membrane fusion. The human furin protein is part of the subtilisin-like serine endoprotease family and recognizes polybasic motifs, having Arg-X-Lys/Arg-Arg (RXK/RR) as a consensus cleavage site [16]. HIV-1 Env possess a highly conserved furin cleavage site at the gp120-gp41 junction (^508^REKR^511^) which is adjacent to the hydrophobic fusion peptide at the gp41 N-terminus, with furin cleavage being essential for viral infectivity [6, 8, 17, 18]. A putative secondary furin cleavage site (^500^KAKR^503^), located a few residues upstream of the primary cleavage site, has been described but its function remains unclear [17, 19].

The functional mature Env trimer is known to sample different conformations ranging from the pre-fusion “closed” metastable conformation (State 1) to the CD4-bound “open” conformation (State 3), transitioning through an intermediate asymmetric conformation (State 2) [20, 21]. Env glycoproteins from primary isolates preferentially adopt the State 1 conformation, which is preferentially recognized by broadly-neutralizing antibodies (bNAbs) [20, 22–24], and can be triggered into downstream conformations by CD4 binding, which exposes highly conserved epitopes targeted by non-neutralizing antibodies (nnAbs) [20, 25, 26]. These nnAbs are rapidly elicited upon infection and vaccination [27–32] and mediate potent Fc-effector functions, including antibody-dependent cellular cytotoxicity (ADCC) [26, 33–38]. The binding of Env to CD4 on the surface of HIV-1-infected cells stabilizes Env in State 2A, which is highly susceptible to nnAbs-mediated ADCC [26, 39, 40]. However, HIV-1 has evolved to prevent the premature adoption of the CD4-bound conformation by downregulating and degrading pre-existing and newly-synthesized CD4 through its accessory proteins Nef and Vpu [26, 35, 41, 42]. Small CD4 mimetic compounds (CD4mc) are being developed to “open up” Env, with the goal of harnessing the potential of nnAbs responses for prevention [31, 32, 38, 43–46] and eradication [36, 40, 47–53] strategies. Another class of Env antagonists known as conformational blockers, which includes the FDA-approved drug Temsavir, prevent Env transitions to downstream conformations by stabilizing Env State 1 [20, 22, 54, 55].

Besides Env-CD4 interaction, there are also structural features of HIV-1 Env that can modulate the sensitivity of HIV-1 to ADCC responses mediated by nnAbs present in plasma from infected individuals. Natural polymorphisms in the Phe43 cavity (notably in CRF01_AE strains) and mutations of conserved residues in the trimer association domain have been shown to modulate Env conformation [25, 56–59] and as a result, the susceptibility of cells infected with these viruses to ADCC responses [51, 60, 61]. Similarly, proteolytic cleavage has been reported to stabilize a “closed” Env conformation [62–65], since mutations in the furin cleavage site resulted in the spontaneous sampling of downstream conformations, including Env State 2A [40, 55, 63]. Here we evaluate the impact of altering Env furin cleavage site on the susceptibility of infected primary CD4+ T cells to ADCC responses mediated by HIV+ plasma.

## 2. Materials and methods

### 2.1 Ethics Statement

Written informed consent was obtained from all study participants [the Montreal Primary HIV Infection Cohort [66, 67] and the Canadian Cohort of HIV Infected Slow Progressors [68–70], and research adhered to the ethical guidelines of CRCHUM and was reviewed and approved by the CRCHUM institutional review board (ethics committee, approval number CE 16.164 - CA). Research adhered to the standards indicated by the Declaration of Helsinki. All participants were adult and provided informed written consent prior to enrolment in accordance with Institutional Review Board approval.

### 2.2 Cell lines and primary cells

293T human embryonic kidney cells (obtained from ATCC) and TZM-bl cells (NIH AIDS Reagent Program) were maintained at 37°C under 5% CO_2_ in Dulbecco’s Modified Eagle Medium (DMEM) (Wisent), supplemented with 5% fetal bovine serum (FBS) (VWR) and 100 U/mL penicillin/streptomycin (Wisent). 293T cells were derived from 293 cells, into which the simian virus 40 T-antigen was inserted. TZM-bl were derived from HeLa cells and were engineered to stably express high levels of human CD4 and CCR5 and to contain the firefly luciferase reporter gene under the control of the HIV-1 promoter [71]. Primary human PBMCs and CD4+ T cells were isolated, activated and cultured as previously described [26]. Briefly, PBMCs were obtained by leukapheresis from six HIV-negative individuals (all males) and CD4+ T lymphocytes were purified from resting PBMCs by negative selection using immunomagnetic beads per the manufacturer’s instructions (StemCell Technologies) and were activated with phytohemagglutinin-L (10 μg/mL) for 48 hours and then maintained in RPMI 1640 complete medium supplemented with rIL-2 (100 U/mL).

### 2.3 Antibodies and sera

The following Abs were used to assess Env conformation at the cell surface: conformation-independent anti-gp120 outer-domain 2G12 (NIH AIDS Reagent Program), broadly-neutralizing antibodies VRC03 (NIH AIDS Reagent Program), PG9 (Polymun), PGT126, PGT151 (IAVI) 10-1074 (kindly provided by Michel Nussenzweig) and VRC34 (kindly provided by John Mascola) as well as non-neutralizing antibodies F240, 19b, 17b, A32, C11 (NIH AIDS Reagent Program). The HIV-IG polyclonal antibody consists of anti-HIV immunoglobulins purified from a pool of plasma from HIV+ asymptomatic donors (NIH AIDS Reagent Program). Goat anti-human and anti-mouse antibodies pre-coupled to Alexa Fluor 647 (Invitrogen) were used as secondary antibodies in flow cytometry experiments. Plasma from HIV-infected individuals were collected, heat-inactivated and conserved at −80 °C until use.

### 2.4 Small molecules

The small-molecule CD4-mimetic compound BNM-III-170 was synthesized as described previously [72]. The HIV-1 attachment inhibitor Temsavir (BMS-626529) was purchased from APExBIO. The compounds were dissolved in dimethyl sulfoxide (DMSO) at a stock concentration of 10 mM and diluted to 50 μM in phosphate-buffered saline (PBS) for cell-surface staining and virus capture assay or in RPMI 1640 complete medium for ADCC assays.

### 2.5 Plasmids and proviral constructs

The vesicular stomatitis virus G (VSV-G)-encoding plasmid was previously described [73]. Transmitted/Founder (T/F) infectious molecular clones (IMCs) of patients CH058 and CH077 were previously reported [74–77]. To generate IMCs encoding for cleavage-deficient Env, two mutations (R508S/R511S) were introduced in the furin cleavage site (^508^REKR^511^) using the QuikChange II XL site-directed mutagenesis protocol (Stratagene). The presence of the desired mutations was determined by automated DNA sequencing.

### 2.6 Radioactive labeling and immunoprecipitation of envelope glycoproteins

3 × 10^5^ 293T cells were transfected by the calcium phosphate method with the different IMCs. One day after transfection, cells were metabolically labeled for 16h with 100 μCi/mL [^35^S]methionine-cysteine ([^35^S] Protein Labeling Mix; Perkin-Elmer) in Dulbecco’s modified Eagle’s medium lacking methionine and cysteine and supplemented with 5% dialyzed fetal bovine serum. Cells were subsequently lysed in RIPA buffer (140 mM NaCl, 8 mM Na_2_HPO_4_, 2 mM NaH_2_PO_4_, 1% NP40, 0.05% sodium dodecyl sulfate (SDS), 1.2 mM sodium deoxycholate). Precipitation of radiolabeled envelope glycoproteins from cell lysates or medium was performed with a mixture of sera from HIV-1-infected individuals in the presence of 50 μl of 10% Protein A-Sepharose (Cytiva) at 4 °C. The precipitated proteins were loaded onto SDS-PAGE gels and analyzed by autoradiography and densitometry to calculate their processing indexes. The processing index is a measure of the conversion of the mutant gp160 Env precursor to mature gp120, relative to that of the wild-type Env trimers. The processing index is calculated with the following formula: processing index = ([total gp120]mutant × [gp160]WT)/([gp160]mutant × [total gp120]WT).

### 2.7 Viral production and infections

To achieve similar levels of infection in primary CD4^+^ T cells among the different IMCs tested, VSV-G-pseudotyped HIV-1 viruses were produced and titrated as previously described [60]. Viruses were then used to infect activated primary CD4+ T cells from healthy HIV-1 negative donors by spin infection at 800 × *g* for 1 h in 96-well plates at 25 °C. To assess viral infectivity, TZM-bl reporter cells were seeded at a density of 2 × 10^4^ cells/well in 96-well luminometer-compatible tissue culture plates (PerkinElmer) 24 h before infection. Normalized amounts of viruses (according to reverse transcriptase activity [78]) in a final volume of 100 μl were then added to the target cells and incubated for 48 h at 37°C. The medium was then removed from each well, and the cells were lysed by the addition of 30 μl of passive lysis buffer (Promega) and one freeze-thaw cycle. An LB 941 TriStar luminometer (Berthold Technologies) was used to measure the luciferase activity of each well after the addition of 100 μl of luciferin buffer (15 mM MgSO_4_, 15 mM KH_2_PO_4_ [pH 7.8], 1 mM ATP, and 1 mM 170 dithiothreitol) and 50 μl of 1 mM d-luciferin potassium salt (Prolume).

### 2.8 Virus capture assay

The HIV-1 virus capture assay was previously reported [79]. Pseudoviral particles were produced by transfecting 2 × 10^6^ 293T cells with pNL4.3 R-E-Luc (NIH AIDS Reagent Program) (3.5 μg), HIV-1_CH058_ (3.5 μg), and VSV-G (1 μg) using standard calcium phosphate method. Forty-eight hours later, virus-containing supernatant were collected, and cell debris were removed by centrifugation (1,500 rpm for 10 min). Anti-Env antibodies were immobilized on white MaxiSorp ELISA plates (Thermo Fisher Scientific) at a concentration of 5 μg/ml in 100 μL of PBS overnight at 4°C. Unbound antibodies were removed by washing twice the plates twice with PBS. Plates were subsequently blocked with 3% bovine serum albumin (BSA) in PBS for 1 h at room temperature. After washing plates twice with PBS, 200 μl of virus-containing supernatants were added to the wells. After 4 to 6 h incubation, virions were removed, and the wells were washed 3 times with PBS. Virus capture by any given antibody was visualized by adding 1 × 10^4^ 293T cells per well in complete DMEM. To measure recombinant virus infectivity, 1 × 10^4^ 293T cells were directly mixed with 100 μl of virus-containing supernatants per well. Forty-eight hours post-infection, cells were lysed by the addition of 30 μl of passive lysis buffer (Promega) and one freeze-thaw cycle. An LB 941 TriStar luminometer (Berthold Technologies) was used to measure the luciferase activity of each well after the addition of 100 μl of luciferin buffer (15 mM MgSO_4_, 15 mM KH_2_PO_4_ [pH 7.8], 1 mM ATP, and 1 mM dithiothreitol) and 50 μl of 1 mM d-luciferin potassium salt (Prolume).

### 2.9 Flow cytometry analysis of cell-surface and intracellular staining

Cell-surface staining of HIV-1-transfected and HIV-1-infected cells was executed as previously described [35, 61]. For transfected cells, we used the standard calcium phosphate method to transfect 7 μg of each IMC into 2 × 10^6^ 293T cells. Binding of cell-surface HIV-1 Env by anti-Env mAbs (5 μg/mL) or HIV+ plasma (1:1000 dilution) was performed at 48h post-transfection. Similarly, cell-surface staining of infected cells was performed at 48h post-infection. After cell-surface staining, transfected cells and infected cells were permeabilized using the Cytofix/Cytoperm Fixation/ Permeabilization Kit (BD Biosciences) and stained intracellularly using PE-conjugated mouse anti-p24 mAb (clone KC57; Beckman Coulter; 1:100 dilution). The percentage of transfected or infected cells (p24^+^) was determined by gating on the living cell population on the basis of a viability dye staining (Aqua Vivid, Thermo Fisher Scientific). Samples were acquired on an LSRII cytometer (BD Biosciences), and data analysis was performed using FlowJo v10.5.3 (Tree Star).

### 2.10 FACS-based ADCC assay

Measurement of ADCC using the FACS-based assay was performed at 48h post-infection as previously described. Briefly, HIV-1-infected primary CD4+ T cells were stained with AquaVivid viability dye and cell proliferation dye (eFluor670; eBioscience) and used as target cells. Autologous PBMC effectors cells, stained with another cellular marker (cell proliferation dye eFluor450; eBioscience), were added at an effector: target ratio of 10:1 in 96-well V-bottom plates (Corning). A 1:1000 final dilution of HIV+ plasma was added to appropriate wells and cells were incubated for 5 min at room temperature. The plates were subsequently centrifuged for 1 min at 300 *x* g, and incubated at 37°C, 5% CO_2_ for 5h before being fixed in a 2% PBS-formaldehyde solution. Samples were acquired on an LSRII cytometer (BD Biosciences) and data analysis was performed using FlowJo v10.5.3 (Tree Star). The percentage of ADCC was calculated with the following formula: (% of p24+ cells in Targets plus Effectors) - (% of p24+ cells in Targets plus Effectors plus sera) / (% of p24+ cells in Targets) by gating on infected lived target cells.

### 2.11 Statistical analysis

Statistics were analyzed using GraphPad Prism version 9.1.0 (GraphPad). Every data set was tested for statistical normality and this information was used to apply the appropriate (parametric or nonparametric) statistical test. P values <0.05 were considered significant; significance values are indicated as * P<0.05, ** P<0.01, *** P<0.001, **** P<0.0001.

## 3. Results

### 3.1 Conformation of HIV-1 uncleaved Env at the surface of infected cells and viral particles

To study the role of the furin cleavage site on Env conformation, we performed mutagenesis on the infectious molecular clones (IMCs) of clade B transmitted/founder (T/F) viruses CH058 and CH077. Envs from both viruses were previously shown to preferentially sample the “closed” State 1 conformation [61]. We introduced substitutions in the primary cleavage site at position 508 and 511 (Figure 1A), to replace the highly conserved arginine residues with serine residues (R508S/R511S; referred as Cl- mutant), a double mutant known to efficiently abrogate furin-dependant Env processing [64, 80–82]. We used protein radioactive labelling of 293T cells transfected with the different IMC constructs followed by Env immunoprecipitation to confirm the effect of the mutations on Env cleavage (Figure 1B-E). As expected, Env glycoproteins expressed from the wild-type (WT) construct were efficiently cleaved while their cleavage-deficient (Cl-) counterpart yielded little to no detectable gp120 in the 293T cell lysates (Figure 1B,D). Although we observed some soluble gp120 in the supernatant of CH058-transfected cells, this was likely due to the presence of second upstream cleavage site, which matched the furin consensus sequence (RAKR). Supernatant of CH077-transfected cells did not contain gp120 consistent with an altered upstream cleavage site (KAKR) (Figure 1A). Of note, two bands of gp160 with distinct molecular weights were observed in cells transfected with Cl- variants, a phenotype previously observed that was linked to a difference in glycosylation [83–85].

**Figure 1.**
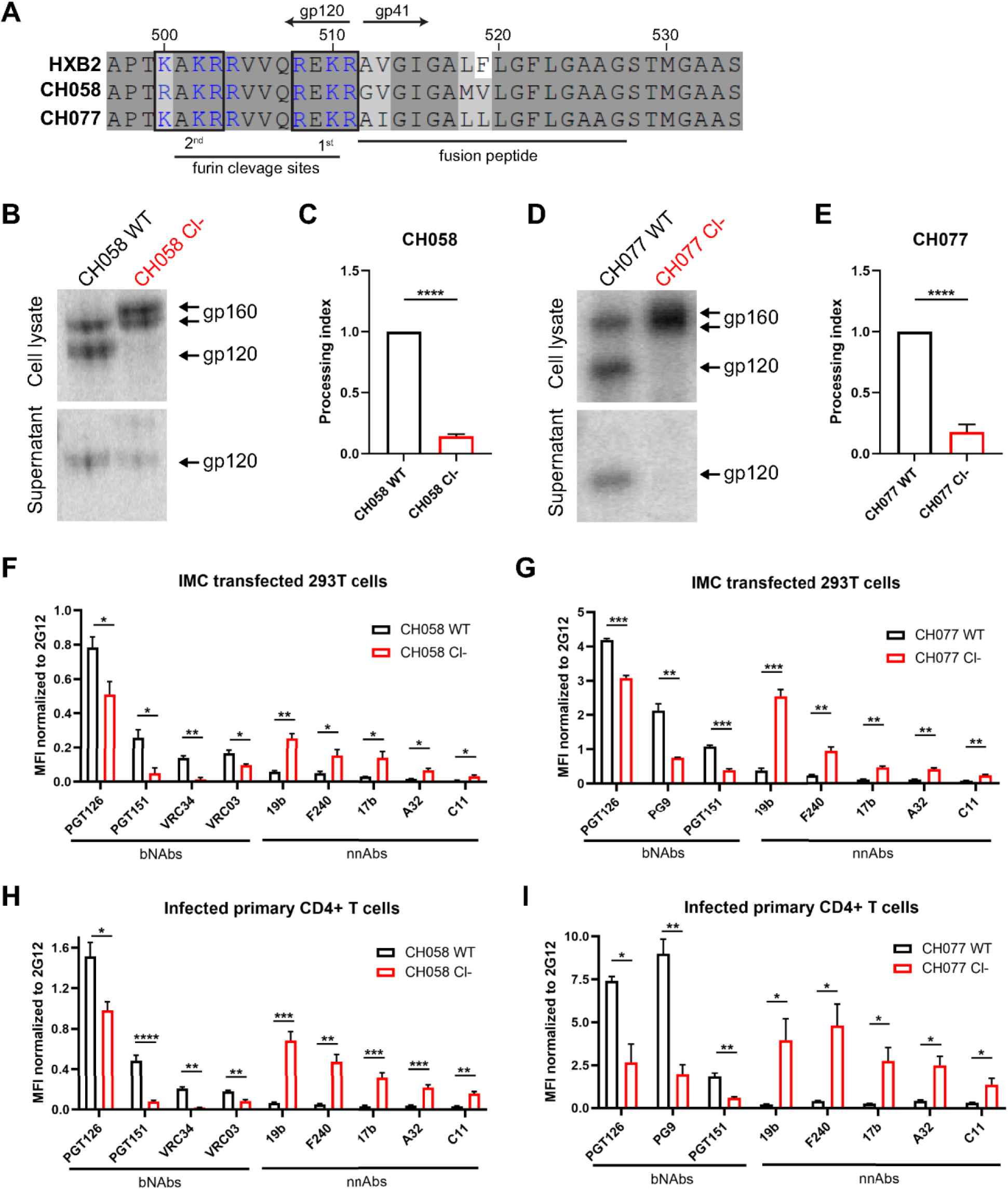
Proteolytic cleavage stabilizes Env in its “closed” conformation. (**A**) Sequence alignment of the HIV-1 Env furin cleavage site region from primary viruses CH058 (GenBank accession number JN944940) and CH077 (GenBank accession number JN944941) with the HXB2 reference strain (GenBank accession number K03455). Putative furin cleavage sequences are highlighted by black boxes. Positively-charged residues (arginine and lysine) are shown in blue. Residue numbering is based on the HXB2 strain. Identical residues are shaded in dark gray, and conserved residues are shaded in light gray. (**B-E**) 293T cells were transfected with primary IMCs (**B-C**) CH058, (**D-E**) CH077 WT or their cleavage-deficient (Cl-) variants and metabolically-labeled with [^35^S]-methionine and [^35^S]-cysteine. (**B,D**) Cell lysates and supernatants were immunoprecipitated with plasma from HIV-1-infected individuals. The precipitated proteins were loaded onto SDS-PAGE gels and analyzed by autoradiography and densitometry to calculate their processing indexes. The processing index is a measure of the conversion of the mutant gp160 Env precursor to mature gp120, relative to that of the wild-type Env trimer. (**C,E**) Shown is the average of processing indexes calculated in 3 independent experiments. (**F-I**) Cell-surface staining of (**F-G**) IMC transfected 293T cells (**H-I**) or primary CD4+ T cells infected with IMCs (**F,H**) CH058 and (**G,I**) CH077 WT or their cleavage-deficient (Cl-) variants using a panel of anti-Env bNAbs (PGT126, PG9, PGT151, VRC34, VRC03) and nnAbs (19b, F240, 17b, A32, C11). Shown are the mean fluorescence intensities (MFI) using the different antibodies normalized to the signal obtained with the conformation-independent 2G12 mAb. MFI values were measured on the transfected or infected (p24+) population for staining obtained in at least 3 independent experiments. Error bars indicate mean ± SEM. Statistical significance was tested using an unpaired t-test (* p < 0.05, ** p < 0.01, *** p < 0.001, **** p < 0.0001).

Subsequently, we evaluated the ability of a panel of bNAbs and nnAbs to recognize the cleaved (WT) and uncleaved (Cl-) Env at the surface of 293T cells. We selected these cells since they don’t express CD4 and it has been well documented that the presence of CD4 affects Env conformation [26, 35, 86]. Cell were transfected with the different IMC constructs and virus-expressing cells were identified using Gag p24 staining (Figure 1F-G). Cell-surface Env expression was normalized using the conformation-independent 2G12 antibody. Cells expressing WT Env were preferentially recognized by the bNAbs preferring the State 1 conformation (PGT126, VRC03, PG9) and recognizing the fusion peptide (PGT151, VRC34) compared to those expressing the respective cleavage site mutants (Figure 1F-G). Conversely, the binding of nnAbs targeting the downstream conformations States 2/3 (19b, F240, 17b) and State 2A (A32, C11) was significantly enhanced on cells expressing uncleaved Env (Figure 1F-G). To confirm this phenotype in a physiologically more relevant culture system, we infected activated primary CD4+ T cells with the different primary IMCs. Of note, all viruses were pseudotyped with the VSV G glycoprotein to normalize the level of infection and to compensate the inability of uncleaved Env to mediate viral fusion. Consistent with the 293T results, productively-infected cells (p24+ CD4_low_) were more efficiently recognized by bNAbs when expressing cleaved Env, and by nnAbs when expressing uncleaved Env (Figure 1H-I). Overall, these results support and extend previous observations indicating that furin cleavage favors the adoption of the native “closed” conformation at the cell surface [40, 65, 84].

We next investigated the effect of furin cleavage on Env conformation at the surface of viral particles, since the viral membrane is known to be enriched in cholesterol, a lipid known to stabilize Env State 1 conformation by interacting with gp41 membrane proximal external region (MPER) [87–89]. Since virions expressing the Env Cl- variants were unable to infect even highly permissive cells, we used a recently developed virus capture assay [79] (Figure 2A). Specifically, we generated luciferase reporter pseudovirions that contained both HIV-1 Env and VSV G glycoproteins, thus allowing captured virions to infect 293T cells in an Env-independent manner (i.e., 293T infection is driven by the incorporated VSV G glycoprotein, Figure 2B). Virions harboring WT Env were captured more efficiently by bNAbs, while virions harboring uncleaved Env were primarily bound by nnAbs (Figure 2C). The recognition of pseudovirions was also assessed using purified anti-HIV-1 immunoglobulins from HIV+ asymptomatic donors (HIV-IG) [90]. Since the vast majority of naturally-elicited antibodies targets Env in its “open” conformation, HIV-IG polyclonal antibodies captured viral particles displaying immature Env in a larger proportion (Figure 2D). HIV-IG specific capture of uncleaved or cleaved Env could be further increased using the small molecule CD4mc BNM-III-170, which stabilizes the CD4-bound conformation (Figure 2D). Alternatively, treatment with the conformational blocker Temsavir decreased the capacity of HIV-IG to capture viral particles bearing Cl- Envs (Figure 2D), in agreement with its capacity to stabilize the “closed” conformation [20, 22, 55]. These results indicate that uncleaved Env can be forced into “open” or “closed” conformations using small molecule Env antagonists.

**Figure 2.**
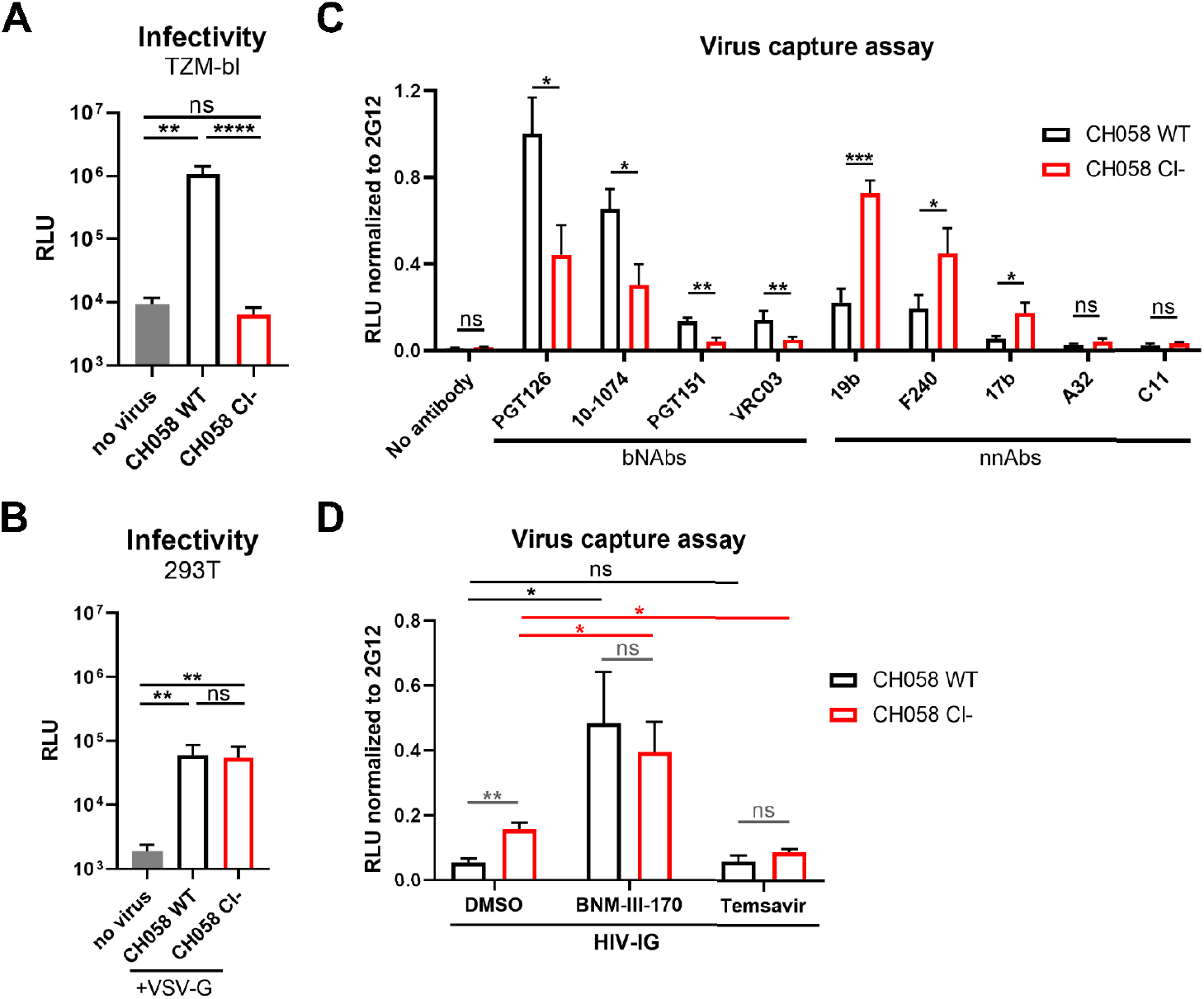
Virions displaying uncleaved Env are better recognized by nnAbs. (**A**) Viral infectivity was assessed by incubating TZM-bl target cells with HIV-1 CH058 virions expressing the wild-type (WT) or cleavage-deficient (Cl-) Env glycoprotein for 48-hours. Viral preparations were normalized according to reverse transcriptase activity. (**B**) VSV-G-pseudotyped viral particles encoding the luciferase gene (Luc+) and bearing HIV-1 CH058 Env wildtype (WT) or its cleavage-deficient mutant (Cl-) were used to infect 293T cells to determine their infectivity in a single-round infection. (**C-D**) These recombinant pseudovirions were further tested for virus capture by (**C**) a panel of anti-Env bNAbs (PGT126, PG9, PGT151, VRC34, VRC03) and nnAbs (19b, F240, 17b, A32, C11) or (**D**) HIV-IG. RLU values obtained using the different antibodies were normalized to the signal obtained with the conformation-independent 2G12 mAb. Data shown are the mean ± SEM from at least three independent experiments. Statistical significance was tested using an unpaired t test or a Mann-Whitney U test based on statistical normality (*, P <0.05; **, P <0.01; ***, P <0.001; ns, nonsignificant).

### 3.2 Impact of HIV-1 Env proteolytic cleavage on ADCC responses mediated by HIV+ plasma

Knowing that alterations in the furin cleavage site increase the exposure of downstream conformations at the surface of infected cells and lentiviral particles, we sought to determine whether the presence of uncleaved Env at the surface of infected cells could also affect ADCC responses mediated by plasma from HIV-1-infected donors. Activated primary CD4+ T cells were infected with WT or cleavage defective CH058 and CH077 and then examined for their susceptibility to ADCC killing following incubation with plasma from 15 different chronically HIV-1-infected individuals. As expected, HIV+ plasma binding was significantly higher on infected cells expressing cleavage-deficient Env compared to WT Env (Figure 3A-B). Moreover, inhibition of Env cleavage led to strong ADCC responses, while WT-infected cells were protected from these responses mediated by HIV+ plasma (Figure 3C-D). Treatment with BNM-III-170 was found to enhance the binding of HIV+ plasma on both WT and Cl- mutant infected cells, consistent with its ability to expose CD4i epitopes. Accordingly, CD4mc addition induced a potent ADCC response against WT-infected cells, but did not further enhance the ADCC response against cells expressing cleavage-deficient Env, suggesting that CD4i epitope exposure by uncleaved Env is sufficient to trigger the elimination of infected cells by ADCC. Conversely, the addition of State 1-stabilizing molecule Temsavir protected Cl- expressing cells from ADCC by decreasing the binding of HIV+ plasma to uncleaved Env (Figure 3A-D). Of note, Temsavir didn’t impact HIV+ plasma mediated ADCC against WT infected cells since they are known to already express the Env in the “closed” conformation [26, 35–37, 40, 60, 61, 86, 91, 92]. Altogether, our results demonstrate the importance for HIV-1 to limit the presence of Env gp160 precursor at the surface of infected cells to evade nnAbs-mediated ADCC responses.

**Figure 3.**
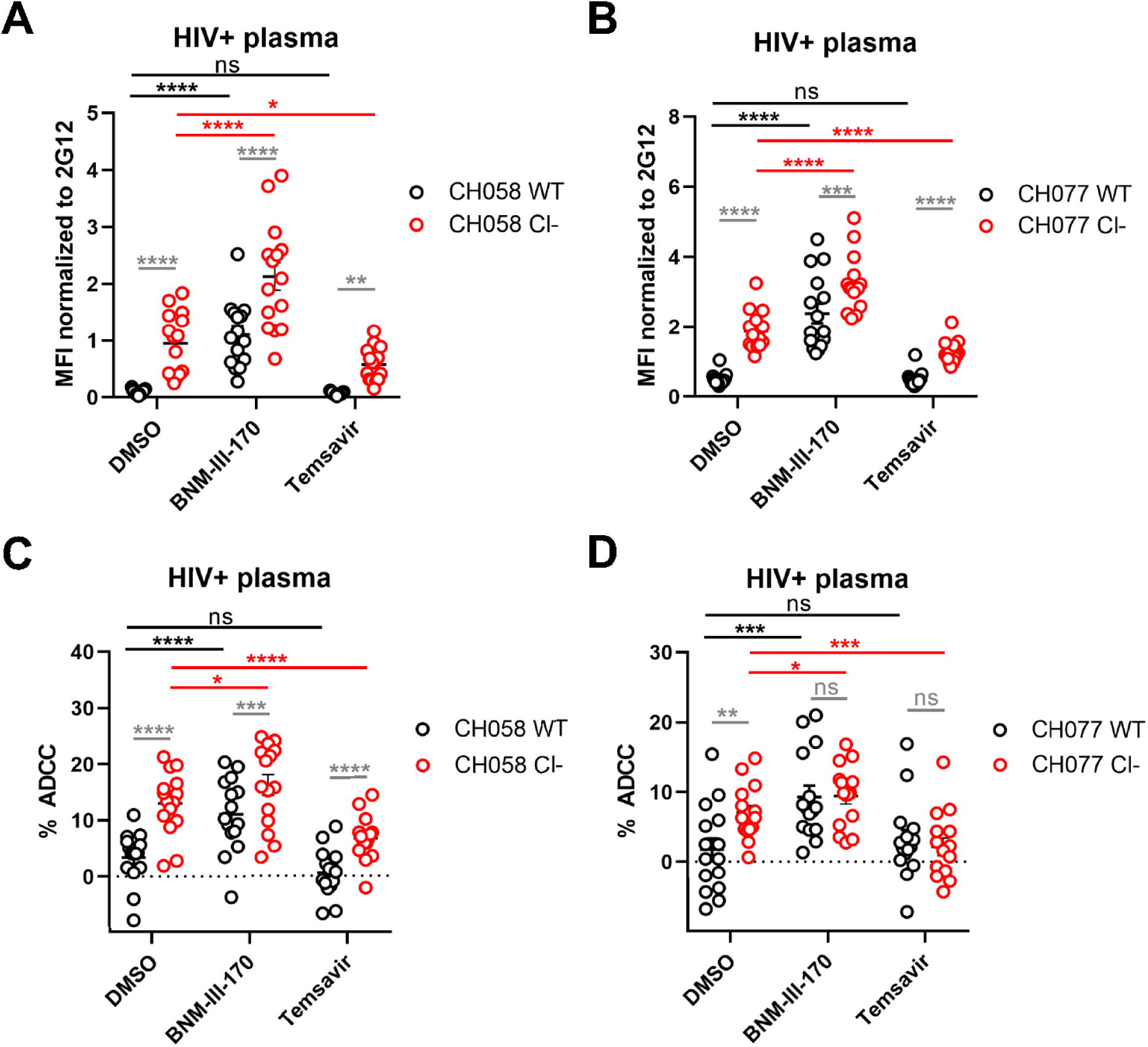
Env cleavage protects HIV-1-infected cells from ADCC mediated by HIV+ plasma. (**A-B**) Cell surface staining of primary CD4+T cells infected with primary HIV-1 viruses (**A**) CH058 and (**B**) CH077 WT or their cleavage-deficient (Cl-) variants using plasma from 15 different HIV-1-infected individuals in the presence of 50 μM of CD4mc BNM-III-170, conformational blocker Temsavir or an equivalent volume of the vehicle (DMSO). The graphs show the MFI obtained on the infected (p24+) cell population. (**C-D**) Primary CD4+ T cells infected with (**C**) CH058 and (**D**) CH077 viruses were also used as target cells, and autologous PBMCs were used as effector cells in a FACS-based ADCC assay. The graphs shown represent the percentages of ADCC mediated by 15 different HIV+ plasma in the presence of 50 μM of CD4mc BNM-III-170, attachment inhibitor Temsavir or an equivalent volume of the vehicle (DMSO). All results were obtained using cells from at least three different donors. Error bars indicate means ± SEM. Statistical significance was tested using a repeated measures one-way ANOVA with a Holm-Sidak post-test (*, P <0.05; **, P <0.01; ***, P <0.001; ****, P <0.0001; ns, nonsignificant).

## 4. Discussion

In this study, we show that uncleaved HIV-1 Env trimers display a conformational flexibility which favors the sampling of downstream “more open” conformations at the surface of infected cells and pseudoviral particles. Cell-surface expression of uncleaved gp160 leads to an efficient recognition of infected cells by non-neutralizing CD4i antibodies naturally-present in plasma from HIV-1-infected individuals and as a consequence, leads to a significantly higher susceptibility to ADCC responses. Conversely, efficient cleavage by endogenous furin allows Env trimers to sample a metastable “closed” conformation (State 1), thus protecting HIV-1-infected cells from ADCC responses mediated by HIV+ plasma. Beyond the well-established role of furin cleavage on viral infectivity, efficient proteolytic cleavage of Env trimers thus appears to allow HIV-1 to evade humoral immune responses. These results are important in the context of recent findings showing that several interferon-inducible cellular antiviral factors affect Env gp160 precursor processing [93–97]. Among them, IFITM proteins impair Env cleavage through a direct interaction with Env, while GBP2 and GBP5 restrict furin protease activity [93, 96]. The antiviral activity of both families of proteins can be overcome by HIV-1 through Env substitutions or by increasing Env expression through the deletion of the accessory Vpu protein, respectively [97–100].

According to the Los Alamos National Laboratory HIV sequence database, very few mutations are naturally found in the furin cleavage site, especially for the basic residues found at position 508, 510 and 511 which are more than 99.7% conserved. Given the importance of an effective Env cleavage to generate infectious viral particles, therapeutic interventions designed to specifically inhibit this proteolytic cleavage could result in a loss in infectivity with a concomitant increase in ADCC responses against infected cells. A recent study has shown that conformational blockers, such as Temsavir, can interfere with proper Env cleavage by reducing its conformational flexibility [63]. Additional drugs inhibiting directly the furin protease activity, including the synthetic peptide Dec-RVKR-CMK and the serine protease inhibitor α1-PDX, are also being investigated, but their *in vivo* efficacy and toxicity remain to be determined [12, 13, 101–105]. If these broad-spectrum inhibitors end up being well tolerated and exhibit good pharmacokinetic properties, they may also be useful as therapeutics against other viral infections, including Influenza A, Ebola, Respiratory syncytial virus (RSV) and SARS-CoV-2, where the acquisition of a furin cleavage site in the respective fusion glycoproteins appears to confer a higher level of infectivity [106–110].

## ACKNOWLEDGMENTS

The authors thank the CRCHUM BSL3 and Flow Cytometry Platforms for technical assistance, Mario Legault from the FRQS AIDS and Infectious Diseases network for cohort coordination and clinical samples. We thank Michel Nussenzweig (The Rockefeller University) for 10-1074 and John Mascola (VRC, NIAID) for VRC34. The graphical abstract was prepared using illustrations from BioRender.com.

## AUTHOR CONTRIBUTIONS

J.P. and A.F. conceived the study. J.P. and A.F. designed experimental approaches. J.P., H.M. and A.F. performed, analyzed, and interpreted the experiments. B.H.H. and A.B.S. supplied novel/unique reagents. J.P. and A.F. wrote the paper. All authors have read, edited, and approved the final manuscript.

## DATA AVAILABILITY

All data are contained within the article.

---

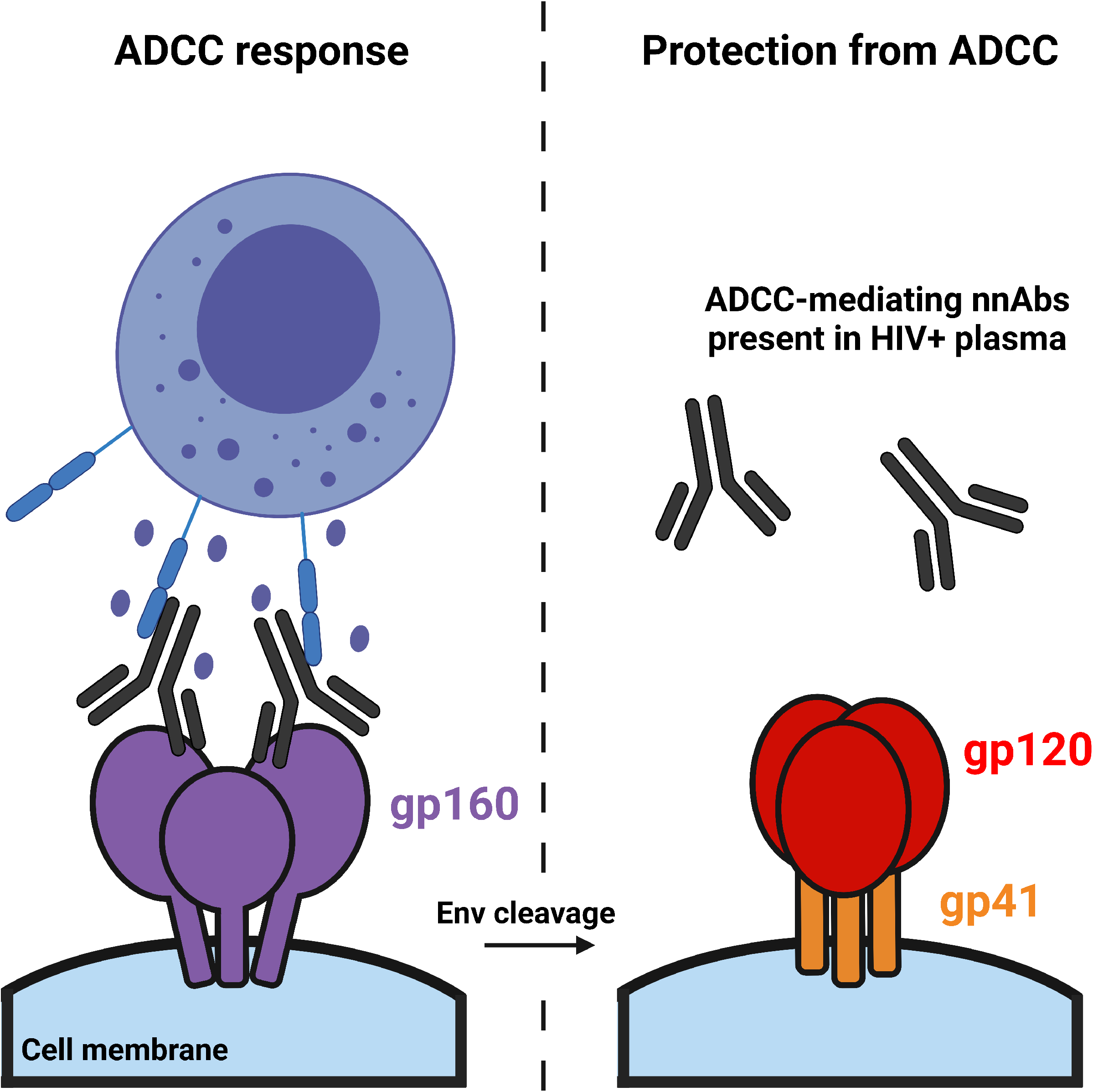

